# Nutrient-responsive midgut turnover is spatially controlled by serotonin signaling

**DOI:** 10.64898/2026.11.03.763885

**Authors:** Nicole Lamichane, Petteri Borgman, Terhi Rintakangas, Kateryna Gaertner, Marius Robciuc, Onur Deniz, Tuuli Kirkinen, Arto Viitanen, Cornélia Biehler, Simon Andersson, Jenny Bärlund, Oona Kinnunen, Pekka Katajisto, Jaakko Mattila, Ville Hietakangas

## Abstract

Barrier epithelia undergo rapid turnover, yet how spatial tissue heterogeneity and environmental cues shape turnover rates remains poorly understood. Here, we show that serotonin signaling spatially regulates homeostatic turnover of the *Drosophila* midgut epithelium in a nutrient-responsive manner. Organ-wide analysis revealed regions of constitutively high and low turnover, as well as areas whose renewal depends on nutrient intake. In rapidly renewing posterior regions, serotonin accumulates in enterocytes undergoing programmed cell death (erebosis) in response to nutrient availability. Serotonin signals through the conserved 5HT7 receptor to promote enteroblast (EB) maturation and migration-associated morphological changes. 5HT7 signaling is both necessary and sufficient for renewal in high-turnover regions, and its sustained disruption leads to epithelial multilayering. Mechanistically, 5HT7 signaling regulates genes involved in metabolic pathways, such as the serine biosynthesis pathway, and in the actin cytoskeleton, which drive EB maturation and morphological change, respectively. Together, our findings establish a signaling mechanism that links nutritional cues to the spatial regulation of barrier epithelium turnover.

## INTRODUCTION

The *Drosophila* midgut is a genetically tractable model to study turnover of barrier epithelium, which is composed of absorptive enterocytes (ECs), their progenitors enteroblasts (EBs) and secretory enteroendocrine (EE) cells, which are renewed by tissue-resident intestinal stem cells (ISCs) (Miguel-Aliaga et al., 2018). Mechanistically, stress-induced epithelial renewal is known to be controlled through signaling mechanisms, such as EGFR-Ras, Hippo and JAK/STAT pathways, that allow damaged epithelia to activate nearby stem cell proliferation and differentiation (Houtz et al., 2017; Jiang and Edgar, 2009b; Jiang et al., 2009; Karpowicz et al., 2010; Osman et al., 2012; Ren et al., 2010). In contrast, the more subtle processes controlling homeostatic midgut turnover remain less resolved. Previous work has shown that the loss of E-cadherin-mediated cell adhesion in dying enterocytes locally triggers stem cell division by activating EGFR signaling, thereby contributing to epithelial cellular homeostasis (Liang et al., 2017). On the other hand, parallel evidence has been presented that the differentiating progenitors, enteroblasts (EBs), are produced in excess, and they undergo epithelial-to-mesenchymal transition, being able to migrate away from their site of origin (Antonello et al., 2015). This implies that additional mechanisms by which dying enterocytes promote the maturation of nearby EBs and control their migration remain to be elucidated.

Midgut size and cellular composition adapt dynamically to nutrient availability. For example, stem cell division is activated by insulin-like signaling in the fed state (O’Brien et al., 2011). Moreover, feeding-induced mTOR complex 1 (mTORC1) activity controls ISC fate determination and differentiation towards the EB/EC lineage (Mattila et al., 2024; Yilmaz et al., 2012). The midgut is regionally heterogeneous with respect to its anatomy and metabolic gene expression, reflecting the compartmentalized nature of digestion and nutrient absorption (Buchon et al., 2013; Marianes and Spradling, 2013). Extensive regional heterogeneity is also observed with respect to nutrient signaling and the consequent regulation of stem cell division and differentiation (Mattila et al., 2024). It remains to be explored, however, whether the regulation of midgut renewal displays similar regional specificity and how nutrient intake is reflected in the regulation of turnover rate.

By conducting an organ-wide analysis of the homeostatic midgut turnover, we observed that the midgut is indeed renewed in a spatially heterogeneous manner. Our data show that the midgut contains regions of high and low constitutive turnover, as well as areas whose turnover is strongly nutrient-dependent. Moreover, we identify serotonin signaling as a key determinant of this process. Serotonin accumulates in ECs undergoing programmed cell death, erebosis (Bergmann, 2022), and its secretion promotes epithelial turnover by signaling through the conserved 5HT7 receptor. Mechanistically, 5HT7 activity facilitates the maturation of EBs by reprogramming the expression of metabolic genes and promotes the migratory properties of EBs through the regulators of the actin cytoskeleton. Collectively, our findings provide evidence for a nutrient-regulated pathway that coordinates spatially distinct tissue turnover rates and facilitates precise local control of cellular replacement to maintain tissue integrity.

## RESULTS

### The turnover of the midgut epithelium is regional and nutrient-dependent

Midgut ECs undergo turnover mediated by stem cell differentiation through the progenitor cell type EB (**Fig 1A**). To analyze how the rate of EC turnover is controlled in midgut regions (**Fig 1B**) and how it responds to changes in nutrient intake, we used regional quantitative analysis by LAM (Viitanen et al., 2021) in combination with lineage tracing using the esgReDDM-system (Antonello et al., 2015). The fluorescent label of esgReDDM allowed us to distinguish newly emerged ECs based on fluorescence and ploidy (**Fig 1C**) and use the proportion of newly emerged ECs out of all ECs as a measure of epithelial turnover. Lineage tracing was activated by transferring the flies to +29°C for 3 days while maintaining them on a nutrient-poor diet (fasted). Subsequently, flies were fed with a chemically defined holidic (Piper et al., 2014) diet (fed) (**Fig 1D**), and the regional turnover rates were analyzed at 0, 4, 7, 14, and 28 days (**Fig 1D, E, S1A-C**). This analysis revealed striking regional heterogeneity in the epithelial turnover and its nutrient responsiveness. Firstly, some midgut regions displayed very low turnover rates: only <30% of ECs located in R1 or the border regions of R3 had undergone turnover at the longest timepoint (28 days) (**Fig 1E, F**), implying that some midgut regions might not undergo complete renewal during the adult lifespan. Secondly, the high turnover regions displayed differential responsiveness to nutrient intake. While the posterior R4 (R4c) displayed a high turnover rate irrespective of nutrient availability, the high turnover of posterior R2 and anterior/middle R4 was only observed in fed animals (**Fig 1E-G**). Collectively, midgut epithelium turnover is highly region-specific and nutrient-responsive.

**Figure 1.**
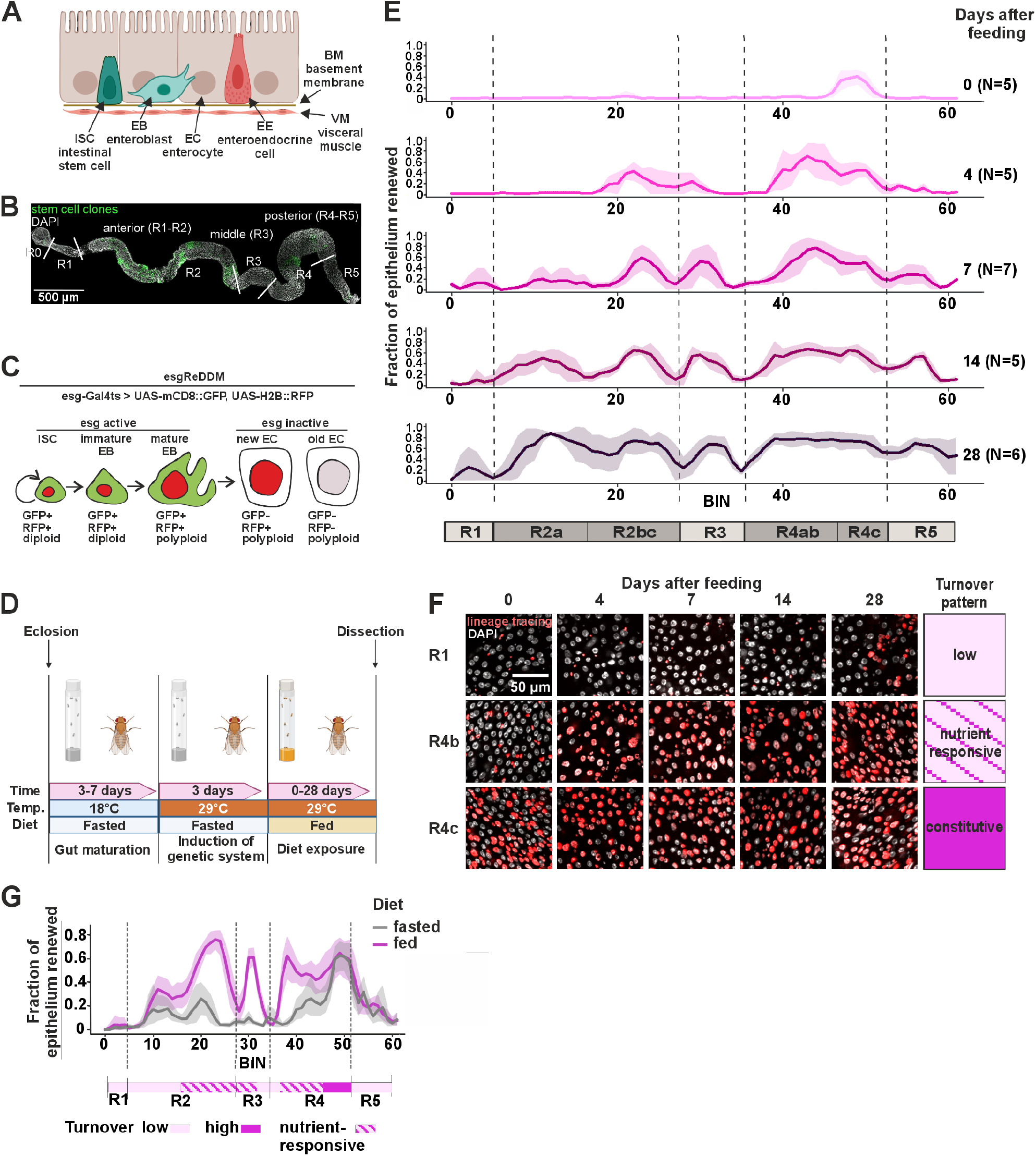
The midgut turnover rate is regional and nutrient-dependent. **A)** Scheme depicting the cellular composition of the *Drosophila* midgut. **B)** Midgut of a fed, mated esgF/O female. Stem cell clones are shown in green (GFP), and nuclei were stained with DAPI (white). **C)** Scheme depicting the esgReDDM system. **D)** Experimental design of the experiments depicted in E-G. Age-matched, mated females (esgReDDM) were matured in fasting conditions at 18°C for 3-7 days, and then shifted to the permissive temperature (29°C) to induce UAS-Gal4-mediated gene expression. After 3 days, flies were transferred to a holidic diet (fed) and maintained on it for the specified period. **E)** Feeding-induced intestinal turnover is regionally heterogeneous. Quantification of intestinal epithelial renewal (esgReDDM: new ECs [polyploid RFP+ and GFP-cells] divided by all ECs [polyploid GFP-cells]) along the midgut 0, 4, 7, 14, and 28 days after feeding of age-matched, mated esgReDDM females. Standard deviation is marked by line shading, the dotted lines indicate the borders of the main intestinal regions. **F)** Representative close-up images from E. DAPI (white) stains nuclei; lineage tracing is shown in red. **G)** Nutrient-responsive turnover is regional. The plot shows the fraction of the intestinal epithelium renewed (new ECs/all ECs) along the midgut, 4 days after feeding vs. fasting, in age-matched, mated esgReDDM females. Standard deviation is marked by line shading.

### Nutrient-regulated enterocyte serotonin promotes midgut turnover

To identify the molecular determinants of nutrient-responsive midgut turnover, we systematically analyzed the impact of single essential amino acids (AAs). Strikingly, feeding of tryptophan (Trp) induced massive turnover specifically in the nutrient-responsive midgut regions, posterior R2 and R4ab (**Fig 2A, B**). Trp can be further metabolized endogenously through the serotonin pathway (**Fig 2C**). To test the possible role of serotonin biosynthesis, we fed esgF/O flies the serotonin precursor 5HTP. Interestingly, feeding 5HTP increased the number of mature EBs, which are not commonly detected under fasting conditions (**Fig 2D, E, Fig 1C**). To gain further insight into the role of serotonin in midgut turnover, we visualized serotonin distribution by immunostaining the midguts with an anti-serotonin antibody (**Fig 2F**). This revealed that serotonin levels showed region-specific distribution, being high in posterior regions and peaking in the R4 (**Fig 2G**).

**Figure 2.**
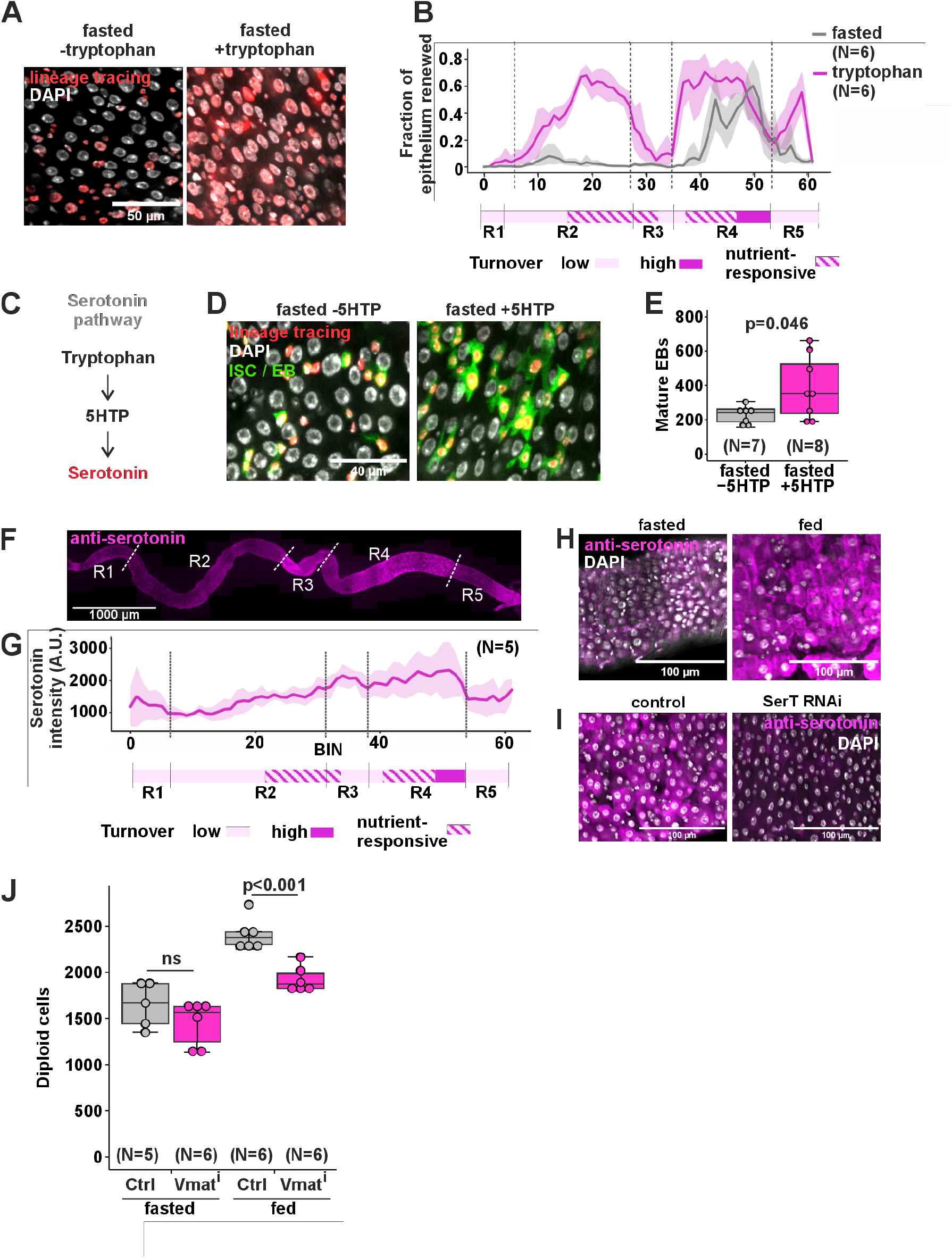
Enterocyte-secreted serotonin regulates intestinal renewal. **A)** Tryptophan feeding promotes intestinal epithelial turnover. Representative images from nutrient-responsive midgut region R4a of age-matched, mated esgReDDM females after 4 days of tryptophan (25 mM) feeding. Lineage tracing (red) was induced with esgReDDM; nuclei are marked with DAPI (white). **B)** Quantification of intestinal epithelial renewal (new ECs/all ECs) along the midgut of age-matched, mated esgReDDM females after 4 days of tryptophan (25 mM) feeding. Standard deviation is marked by line shading. **C)** A scheme depicting the serotonin synthesis pathway. **D)** Dietary serotonin increases stem cell activity. Representative images from midgut region R4a of age-matched, mated esgReDDM females after 4 days of feeding with serotonin-precursor 5HTP (7.5 mM). Lineage tracing (RFP) was induced with esgReDDM, GFP marks ISCs and EBs, and nuclei are marked with DAPI (white). **E)** Dietary serotonin increases EB maturation. Quantification of mature EBs of (D) from posterior midguts. **F)** Serotonin content is highest in the posterior midgut. Representative image of a midgut from a fed, age-matched, mated esgF/O female immunostained with anti-serotonin (magenta). Dotted white lines mark borders between the main midgut regions. **G)** Quantification of serotonin content along the midgut from the experiment depicted in G. Standard deviation is marked by line shading. **H)** Midgut serotonin levels are diet-dependent. Representative image of the midgut region R4c from age-matched, mated esgF/O females after 4 days of feeding or fasting, immunostained with anti-serotonin (magenta). Nuclei are marked with DAPI (white). **I)** Serotonin accumulation in ECs is dependent on serotonin transporter SerT. Representative images of the midgut region R4c from fed, age-matched, mated Myo-GAL4 > luc RNAi or SerT RNAi females immunostained with anti-serotonin (magenta). Nuclei are marked with DAPI (white). **J)** Preventing EC serotonin secretion leads to a decrease in diploid cell numbers (mostly ISCs and EBs). Quantification of diploid cell numbers in the posterior midgut of fed, age-matched, mated Myo-GAL4 > luc RNAi or Vmat RNAi females. P-values in (**E**) were calculated with unpaired t test with Welch correction. P-values in (**J**) were calculated with a two-way ANOVA followed by Tukey’s HSD post-hoc test for pairwise comparisons. N = biological replicates. In box plots, the center line represents the median, the box boundaries represent the 25th and 75th percentiles, and the whiskers extend to 1.5 X IQR from the 25th and 75th percentiles. Individual dots represent separate biological replicates.

This led us to ask whether nutrition regulates serotonin levels in the midgut. Indeed, compared to fed animals, midguts of fasted animals displayed strongly reduced intracellular serotonin levels in R4 (**Fig 2H)**, highlighting the nutrient regulation of midgut serotonin levels. We then wondered whether ECs produce serotonin themselves or whether they rely on serotonin uptake. To test this, we knocked down the serotonin transporter gene SerT specifically in the ECs (Myo1A-GAL4). Knockdown of SerT led to a marked reduction in intracellular serotonin levels in R4 ECs in fed animals (**Fig 2I**), implying that ECs take up extracellular serotonin via SerT.

The regional correlation between EC intracellular serotonin (**Fig 2F**) and the turnover rate (**Fig 1G**) led us to hypothesize that ECs might use serotonin to signal to surrounding progenitors about the need for turnover. To test this, we knocked down Vmat, a monoamine transporter that loads serotonin into vesicles for exocytosis, in the ECs. Interestingly, inhibition of EC serotonin secretion led to a strong nutrient-dependent reduction in the diploid cell population (mostly stem cells and early EBs) in the posterior regions, consistent with nutrient regulation of midgut serotonin levels (**Fig 2J**). In addition, inhibiting EC serotonin uptake by SerT knockdown also reduced the number of diploid cells in the midgut (**Fig S2A)**. Taken together, our results are consistent with a model in which ECs take up and accumulate serotonin in a nutrient-dependent manner and then secrete it to nearby progenitor cells to increase epithelial turnover.

### Serotonin accumulates in ECs undergoing programmed cell death

Homeostatic epithelial turnover requires coordination between EC elimination and differentiation. It was recently demonstrated that midgut ECs undergo an atypical form of programmed cell death, termed erebosis, characterized by large, flattened nuclei, loss of GFP fluorescence and high expression of angiotensin-converting enzyme (ANCE) (Ciesielski et al., 2022). To explore the potential role of erebosis in nutrient-responsive turnover, we analyzed the distribution of erebotic markers with respect to turnover. Firstly, local analysis of the high-turnover region R4b revealed an inverse correlation between ANCE-positive erebotic cells and GFP-marked newly emerged ECs (**Fig 3A**), consistent with the replacement of erebotic ECs by new ECs. Secondly, we observed that the number of ANCE-positive erebotic cells was much higher in the midguts of fed animals, compared to those of fasted animals (**Fig 3B, C**). Moreover, feeding serotonin precursor to fasted animals strongly increased the number of ANCE positive cells (**Fig 3D, E**), demonstrating a causal relationship between serotonin availability and erebosis. To directly compare intracellular serotonin levels with erebosis, we analyzed the erebosis-associated loss of GFP fluorescence (Ciesielski et al., 2022), as simultaneous use of ANCE and serotonin antibodies was not feasible (both derived from the same species). Indeed, ECs with high serotonin content typically lost most of their cytoplasmic GFP-fluorescence (**Fig 3F, G**), consistent with the idea that ECs with particularly high serotonin content are undergoing erebosis.

**Figure 3.**
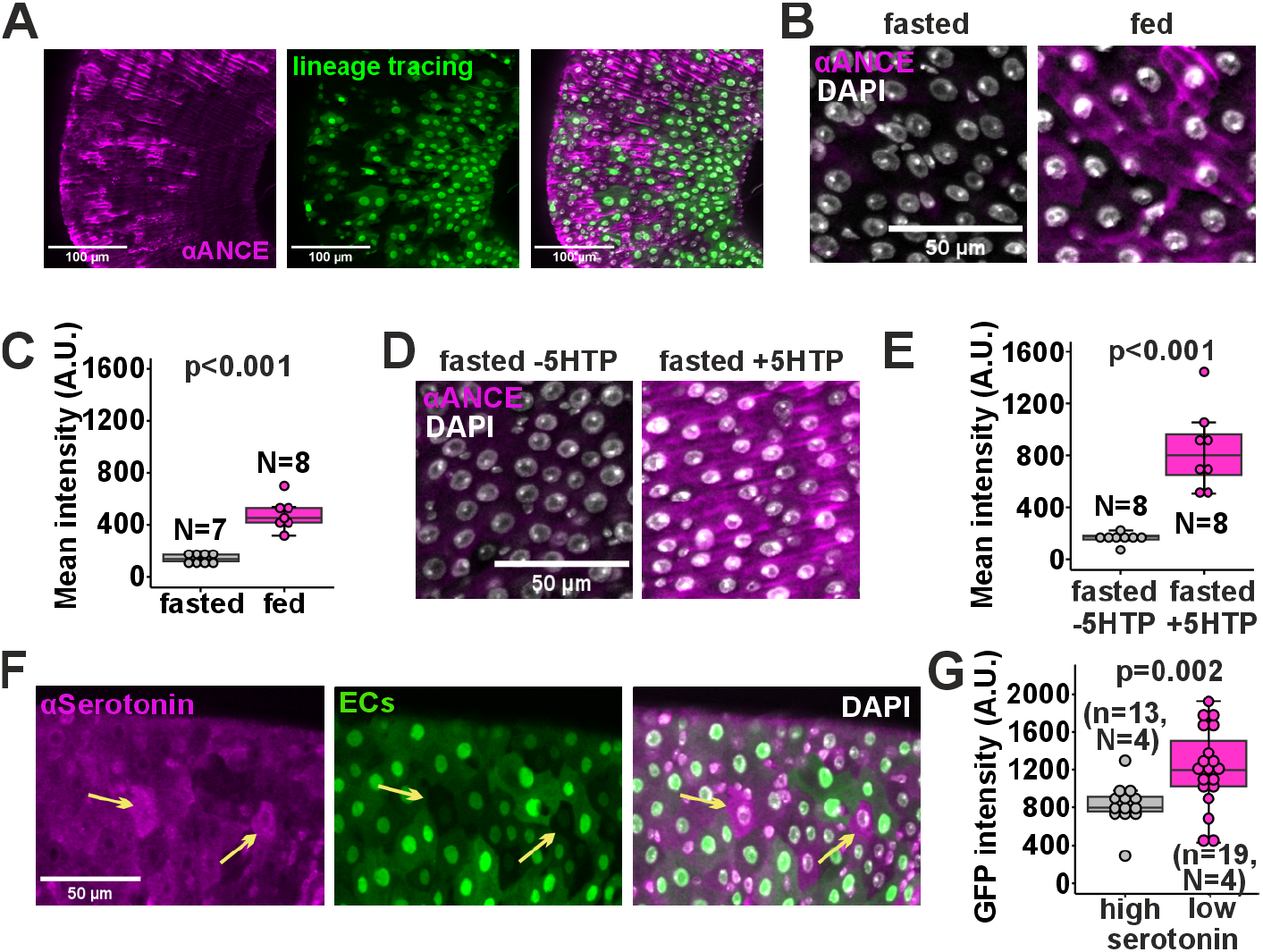
Serotonin accumulates to ECs undergoing erebosis. **A)** Midgut regions exhibit heterogeneous turnover. Representative images of the midgut region R4b from a fed esgF/O female showing that patches with newly generated ECs (GFP+ cells) lack the erebosis marker anti-ANCE (magenta), whereas areas with old ECs contain erebotic ECs. **B & C)** Feeding increases the number of erebotic cells. **B)** Representative images of the midgut region R4b from fasted and fed age-matched, mated esgF/O females. Erebotic cells are visualized by immunostaining with anti-ANCE antibodies (magenta). Nuclei are marked with DAPI (white). **C)** Quantification of the anti-ANCE mean intensity from the experiment depicted in B. **D & E)** Dietary serotonin induces erebosis. **D)** Representative images of the midgut region R4b from fasted, age-matched, mated esgF/O females fed with serotonin precursor 5HTP (7.5 mM), immunostained with erebosis marker anti-ANCE (magenta). Nuclei are marked with DAPI (white). **E)** Quantification of the anti-ANCE mean intensity from the experiment depicted in D. **F & G)** ECs with high serotonin content show erebotic characteristics. **F)** Representative image of the midgut region R4b from a mated, Myo-Gal4 > UAS-GFP female. ECs are labeled with GFP, serotonin is immunostained with anti-serotonin (magenta), and nuclei are marked with DAPI (white). Arrows point to ECs with high serotonin/low GFP signal. **G)** Quantification of mean GFP intensity from ECs of high and low serotonin content from the experiment depicted in F. P-values in (**C, E, G**) were calculated with unpaired t test with Welch correction. N = biological replicates, n = technical replicates. In box plots, the center line represents the median, the box boundaries represent the 25th and 75th percentiles, and the whiskers extend to 1.5 X IQR from the 25th and 75th percentiles. Individual dots represent separate biological replicates, except in (G) where they represent separate technical replicates (cells).

### 5-HT7 is necessary for progenitor differentiation and epithelial turnover

How do stem and progenitor cells sense the serotonin secreted by ECs? Extracellular serotonin mediates signaling through conserved serotonin receptors, of which 5 subtypes exist in *Drosophila*: 5HT1A, 5HT1B, 5HT2A, 5HT2B, and 5HT7. To investigate their involvement in regulating the midgut response to serotonin, we systematically knocked down each receptor subtype in stem cells and progenitors and fed the flies 5HTP to test stem cells’ ability to respond to dietary serotonin precursor (**Fig S3A**). Interestingly, knockdown of one of these receptors, 5HT7, blunted the intestinal stem cell activation normally seen upon 5HTP feeding (**Fig S3A**). Next, we wanted to test whether 5HT7 is also involved in homeostatic intestinal turnover in the absence of ectopic serotonin. Strikingly, loss of 5HT7 in stem and progenitor cells led to an almost complete loss of EC turnover (**Fig 4A, B**), accompanied by a reduction in the number of mature EBs (**Fig S3B**). 5HT7 depletion by crRNA-mediated Cas12a excision phenocopied the decrease in stem cell activity seen with 5HT7 knockdown (**Fig S3C, D**). Notably, 5HT7 expression was necessary for EB/EC differentiation in both the nutrient-responsive regions as well as in the constitutive high turnover region (**Fig 4C**). 5HT7 is conserved in mammals, but its role in regulating intestinal stem cells remains unknown. To test this, we treated intestinal mouse organoids with 5HT7 agonist LP-211. Interestingly, 5HT7 activation significantly increased the number of buds in the organoids (**Fig 4D, E**). This indicates that, although the cell death under homeostatic turnover in mammalian intestines occurs in a separate tissue compartment, 5HT7 signaling can activate intestinal stem cells or progenitors also in mammals.

**Figure 4.**
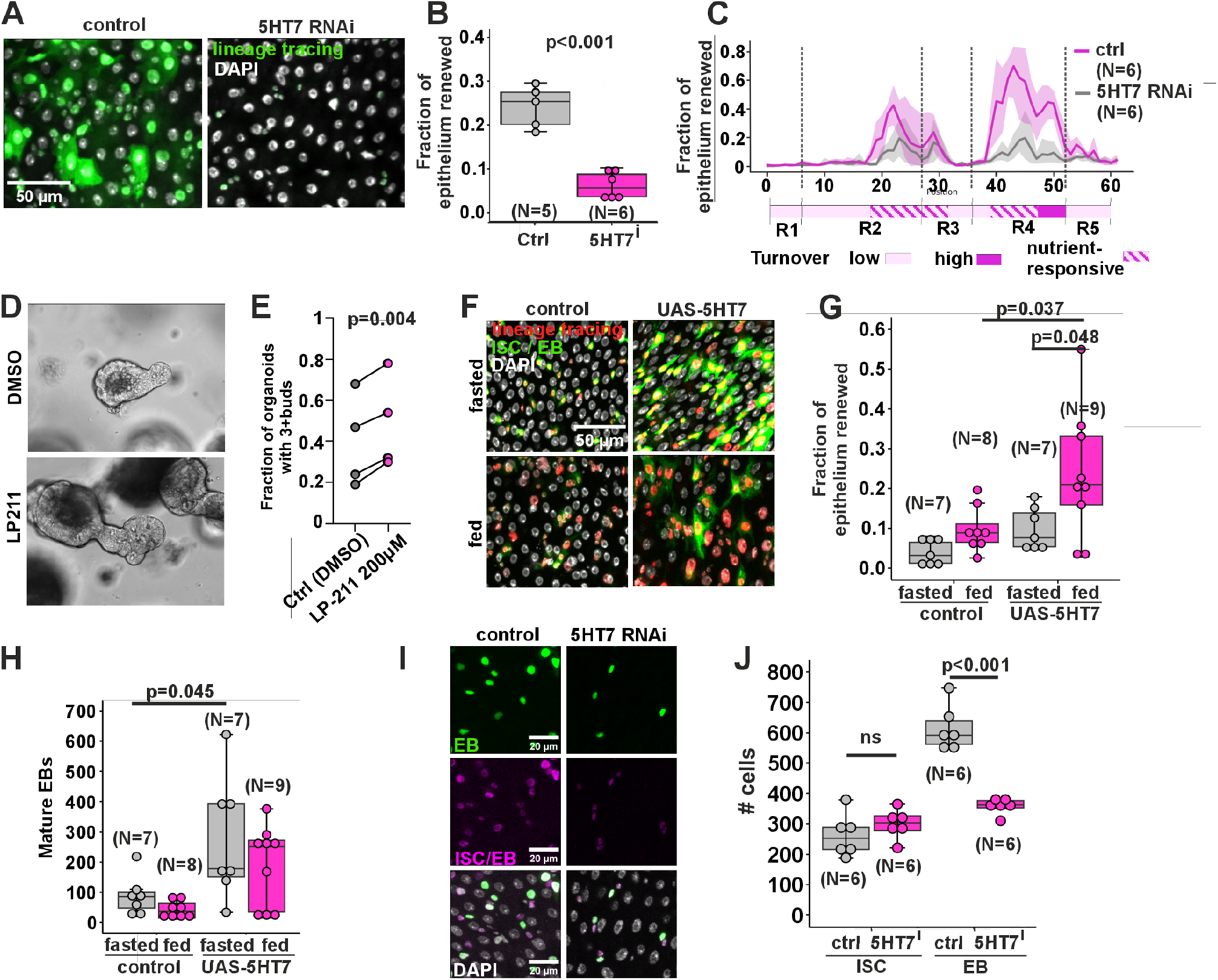
Serotonin receptor 5-HT7 is necessary for progenitor differentiation and epithelial turnover. **A & B)** 5HT7 is needed in ISCs and EBs for intestinal turnover. **A)** Representative images of the midgut region R4c from fed, age-matched, mated esgF/O > luc RNAi or 5HT7 RNAi females. DAPI (white) stains nuclei; lineage tracing is shown in green. **B)** Quantification of intestinal epithelial renewal (new ECs / all ECs) from posterior midguts of fed, age-matched, mated esgReDDM > luc RNAi or 5HT7 RNAi females. **C)** Quantification of intestinal epithelial renewal (new ECs / all ECs) along the midgut of fed, age-matched, mated esgReDDM > luc RNAi or 5HT7 RNAi females. Standard deviation is marked by line shading, the dotted lines indicate the borders of the main intestinal regions. **D & E)** 5HT7 activation leads to increased bud formation in mouse organoids. **D)** Representative bright-field images of mouse organoids after treatment with vehicle (DMSO) or 5HT7 agonist LP-211 (200 µM). **E)** Quantification of the fraction of organoids that developed at least 3 buds from the experiment depicted in D. **F-H)** Overexpression of 5HT7 increases midgut turnover and EB maturation. **F)** Representative images of the posterior midguts from fed and fasted, age-matched, mated esgReDDM > attP40 control or UAS-5HT7 females. DAPI (white) stains nuclei; lineage tracing is shown in red and ISCs/EBs are marked with GFP (green). **G)** Quantification of intestinal epithelial renewal (new ECs / all ECs) of (F) from nutrient-responsive regions R4a-R4b. **H)** Quantification of mature EBs of (F) from posterior midguts.. **I-L)** 5HT7 regulates EB numbers. **I)** Representative images of ISCs (CFP+ [magenta] and GFP-[green]) and EBs (CFP+ and GFP+) from posterior midguts of fed, age-matched, mated Cfs-GAL4 > luc RNAi or 5HT7 RNAi females. **J)** Quantification of ISC and EB numbers from the experiment depicted in I. P-values in (**B, J**) were calculated with unpaired t test with Welch correction. P-values in (**E**) were calculated with a paired t test. P-values in (**G, H**) were calculated with a two-way ANOVA followed by Tukey’s HSD post-hoc test for pairwise comparisons. N = biological replicates. In box plots, the center line represents the median, the box boundaries represent the 25th and 75th percentiles, and the whiskers extend to 1.5 X IQR from the 25th and 75th percentiles. Individual dots represent separate biological replicates.

Since the loss of 5HT7 led to reduced turnover, we wanted to test the consequences of overactive serotonin signaling by using 5HT7 overexpression with the esgReDDM driver. Consistent with loss-of-function data, 5HT7 overexpression led to a striking increase in EC turnover in the posterior midgut of fed animals (**Fig 4F, G**). Is high serotonin signaling sufficient to override the inhibition of EC turnover during fasting? 5HT7 overexpression strikingly increased the number of mature EBs, but was insufficient to promote EC differentiation and turnover (**Fig 4F-H**), similar to the phenotype observed with 5HTP feeding of fasted animals (**Fig 2D**). To further investigate the role of 5HT7 in controlling ISCs and EBs, we used the Cell Fate Sensor (Cfs-GAL4), which employs fluorescent labels to distinguish ISCs and EBs (Martin et al., 2018). Using this system, we observed that knockdown of 5HT7 had no impact on steady-state ISC numbers but significantly reduced the number of EBs (**Fig 4I, J**). Collectively, our data show that signaling through the 5HT7 receptor is both necessary and sufficient to promote progenitor maturation and epithelial turnover in the posterior midgut.

### 5-HT7 signaling controls maturation-promoting metabolism in the progenitor cells

To elucidate the molecular mechanisms by which 5HT7 regulates stem and progenitor cells, we performed RNA sequencing on control, 5HT7-deficient, and 5HT7-overexpressing ISCs and EBs, isolated by FACS (Dutta et al., 2013). Principal component analysis of the RNA-seq data showed that cells with 5HT7 RNAi and those overexpressing 5HT7 clustered separately from each other and from the control (**Fig 5A**), highlighting the significant impact of serotonin signaling on gene expression. Gene set enrichment analysis (GSEA) revealed that genes involved in metabolic pathways were differentially expressed between the control and 5HT7-knockdown groups (**Fig 5B**). For example, 5HT7-regulated metabolic genes include *glutamine-fructose-6-phosphate transaminase 1* (*gfat1*) and *astray* (*aay*) (**Fig 5C-F**), which encode enzymes of the hexosamine and serine biosynthesis pathways, respectively (**Fig 5G**). In addition, genes involved in glycolysis and gluconeogenesis were deregulated upon 5HT7 knockdown (**Fig S4**).

**Figure 5.**
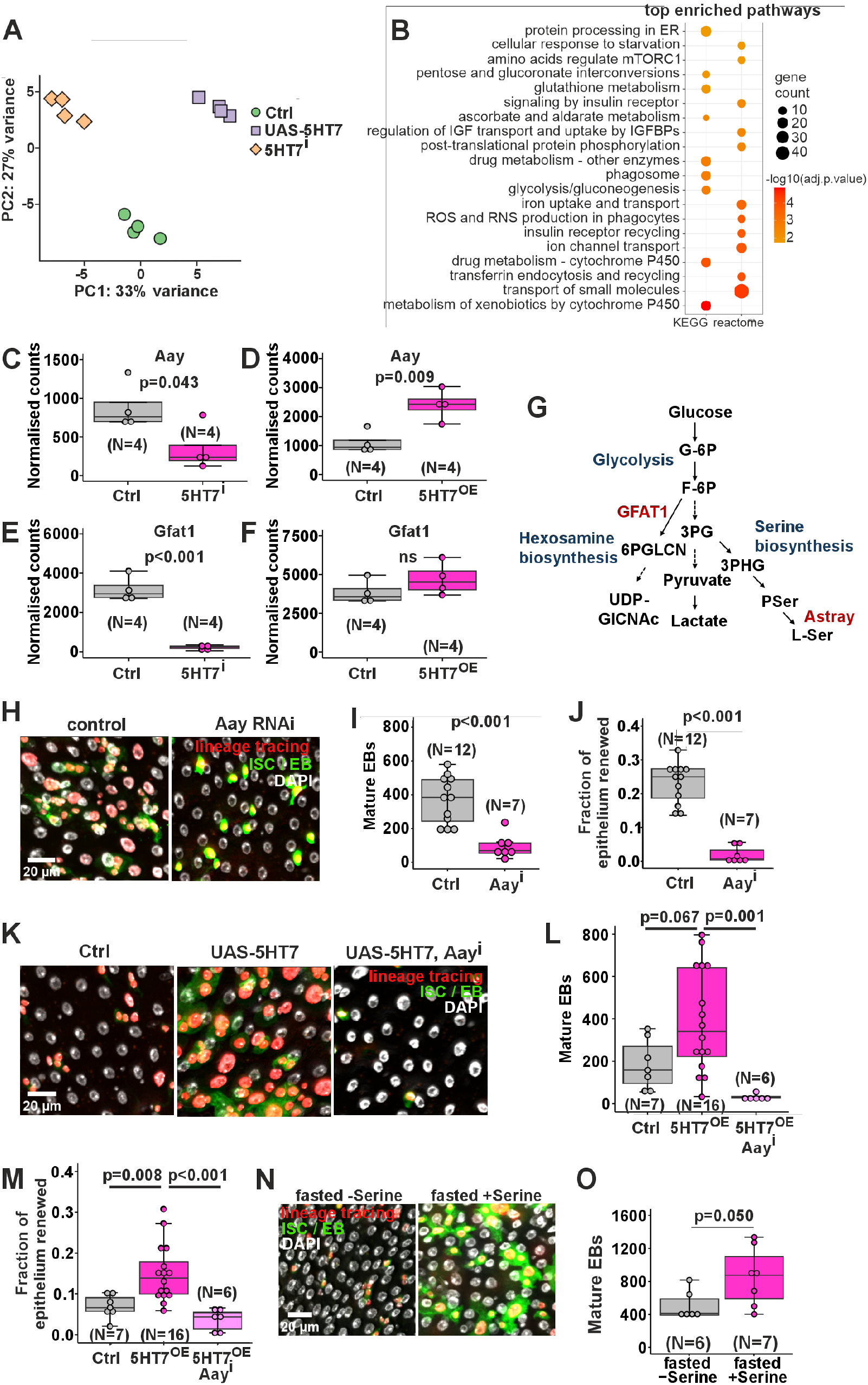
5HT7 regulates metabolic gene expression necessary for EB differentiation. **A-F)** 5HT7 affects gene expression in ISCs and EBs. **A)** Principal component analysis of RNA-sequencing data of FACS-isolated ISCs and EBs of the posterior midgut of fed, age-matched, mated esg-GAL4 > Oregon R (control), 5HT7 RNAi, or UAS-5HT7 females. **B)** Gene set enrichment analysis showing differentially regulated pathways between control and 5HT7-knockdown samples. **C-D)** 5HT7 regulates *aay* expression. Quantifications of normalized mRNA expression from the RNA sequencing experiment depicted in A. **E-F)** 5HT7 regulates *gfat1* expression. Quantifications of normalized mRNA expression from the RNA sequencing experiment depicted in A. **G)** A scheme of glucose metabolism highlighting the role of Aay and Gfat1 in Serine and Hexosamine biosynthesis, respectively. **H-J)** Aay is necessary for EB maturation. **H)** Representative images of posterior midguts from fed, age-matched, mated esgReDDM > luc RNAi or Aay RNAi females. DAPI (white) stains nuclei; lineage tracing is shown in red and ISCs/EBs are marked with GFP (green). **I)** Quantification of mature EBs (polyploid RFP+ and GFP+ cells) from the experiment depicted in F. **J)** Quantification of intestinal epithelial renewal (new ECs/all ECs) of the posterior midguts from the experiment depicted in F. **K-M)** Aay is necessary for 5HT7-dependent EB maturation and epithelial turnover. **K)** Representative images of posterior midguts of fed, age-matched, mated esgReDDM > attP40 control, UAS-5HT7 or UAS-5HT7; Aay RNAi females. DAPI (white) stains nuclei; lineage tracing is shown in red and ISCs/EBs are marked with GFP (green). Quantification of mature EBs (**L**) and intestinal epithelial renewal (new ECs/all ECs) in nutrient-responsive regions R4ab (**M**) of (K**). N & O)** Dietary serine increases EB maturation. **N)** Representative images of the midgut region R4c from fasted, age-matched, mated esgReDDM females fed with 25 mM serine. DAPI (white) stains nuclei; lineage tracing is shown in red and ISCs/EBs are marked with GFP (green). **O)** Quantification of mature EBs (polyploid RFP+ and GFP+ cells) in the posterior midgut from the experiment depicted in K. Adjusted P values in (**B**) were calculated by a one-sided hypergeometric test for over-representation analysis and adjusted using the Benjamini–Hochberg method. Adjusted *P* values in (**C-F**) were obtained in differential expression analysis of RNA-Seq with Benjamini–Hochberg correction. Normalised counts were obtained from the DESeq2 object of the respective comparison; small differences in control values between panels reflect comparison-specific size factor estimation and not different samples. P-values in (**I, J, O**) were calculated with unpaired t test with Welch correction. P-values in (**L, M**) were calculated with a one-way ANOVA followed by Tukey’s HSD post-hoc test for pairwise comparisons. N = biological replicates. In box plots, the center line represents the median, the box boundaries represent the 25th and 75th percentiles, and the whiskers extend to 1.5 X IQR from the 25th and 75th percentiles. Individual dots represent separate biological replicates.

While the role of the hexosamine pathway in promoting ISC activity has already been established (Mattila et al., 2018), we explored further the potential role of serine biosynthesis. Indeed, knockdown of Aay, a cytosolic L-phosphoserine phosphatase, prominently inhibited progenitor cell formation and epithelial turnover (**Fig 5H-J, S3E, F**). Moreover, genetic epistasis experiments demonstrated that knockdown of Aay was sufficient to suppress the 5HT7-mediated increase in progenitor cell numbers (**Fig 5K, L**) and epithelial turnover (**Fig 5K, M**). Further supporting the instructive role of serine metabolism in ISC maturation, we observed that serine feeding was sufficient to promote EB formation under fasted conditions (**Figure 5N, O**). In sum, 5HT7 activity controls the metabolic state of stem cells and progenitors, activating biosynthetic pathways, including serine biosynthesis, that promote ISC maturation and EC turnover.

### Serotonin signaling regulates progenitor morphology and migration to facilitate epithelial turnover

Serotonin-positive erebotic ECs were typically surrounded by progenitors in close contact (**Fig 6A**). Moreover, while 5HT7 knockdown acutely inhibits EB maturation, over longer time periods (14 days), 5HT7-deficient cells were able to mature, but failed to properly integrate into the epithelium, leading to multilayering (**Fig 6B, C**). These observations suggest that serotonin may serve as a signal that facilitates progenitor migration towards the dying enterocyte. This hypothesis was supported by the finding that 5HT7 signaling prominently influenced the morphology of the EBs. While wild-type EBs undergo a morphological change displaying cellular extensions, depletion of 5HT7 expression prevented this transition, resulting in round EB morphology (**Fig 6D, E**). Conversely, EBs overexpressing 5HT7 displayed exceptionally large cellular extensions (**Fig 6F, G**). Consistent with these morphological observations, EBs overexpressing 5HT7 were highly mobile, as visualized by ex vivo live imaging of midguts (Video S1, S2).

**Figure 6.**
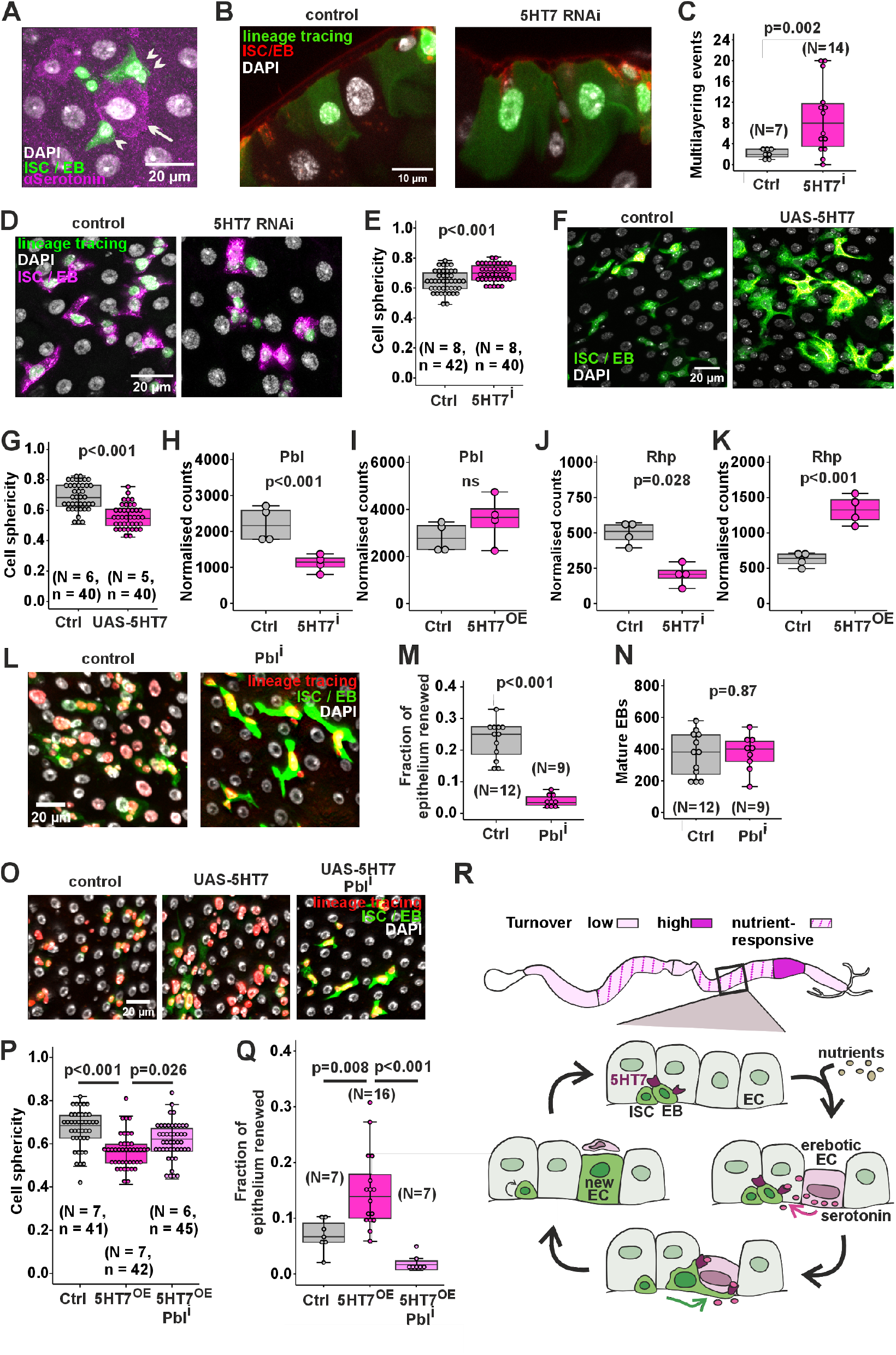
Serotonin signaling regulates progenitor morphology and migration. **A)** Serotonin-positive erebotic ECs (arrow) are surrounded by progenitors (arrowheads) in close contact. Representative image of midgut region R4b from fed, mated esgF/O female. DAPI (white) stains nuclei; anti-serotonin staining is shown in magenta and ISCs/EBs are marked with GFP (green). **B & C)** 5HT7 is necessary for epithelial integration of ECs. **B)** Representative images of midgut region R4c of age-matched mated esgReDDM > luc RNAi or 5HT7 RNAi females after 14 days of feeding. Progenitors are visualized by anti-HRP immunostaining (red), DAPI (white) stains nuclei and lineage tracing is shown in green. **C)** Quantification of multilayering events from the experiment depicted in F. Multilayering events are defined as new ECs differentiating on top of old ECs. **D-G)** 5HT7 regulates EB maturation and morphology. **D)** Representative image of midgut R4b region from fed, age-matched, mated esgF/O > luc RNAi or 5HT7 RNAi females. Progenitors are visualized by anti-HRP immunostaining (magenta), DAPI (white) stains nuclei and lineage tracing is shown in green. **E)** Quantification of mature EB sphericity from the experiment depicted in D. Mature EBs were identified based on their polyploidy and anti-HRP immunostaining. **F)** Representative image of midgut R4b region from fed, age-matched, mated esgReDDM > attP40 control or UAS-5HT7 females. DAPI (white) stains nuclei and ISCs / EBs are marked with GFP (green). **G)** Quantification of mature EB sphericity of (F). **H-I)** 5HT7 regulates *pbl* expression. Quantifications of normalized mRNA expression from the RNA sequencing experiment depicted in Figure 4A. **J-K)** 5HT7 regulates *rhp* expression. Quantifications of normalized mRNA expression from the RNA sequencing experiment depicted in Figure 4A. **L-N**) Pbl is necessary for epithelial renewal but not for EB maturation. **L)** Representative images of the posterior midguts from fed, age-matched, mated esgReDDM > luc RNAi or pbl RNAi females. DAPI (white) stains nuclei, lineage tracing is marked with RFP (red) and ISCs / EBs are marked with GFP (green). **M)** Quantification of intestinal epithelial renewal (new ECs/all ECs) of the posterior midguts of (L). **N)** Quantification of mature EBs (polyploid RFP+ and GFP+ cells) of (L). **O-Q)** Pbl is necessary for 5HT7-induced EB maturation and epithelial renewal. **O)** Representative images of fed, age-matched, mated esgReDDM > attP40 control, UAS-5HT7 or UAS-5HT7; Pbl RNAi females. DAPI (white) stains nuclei, lineage tracing is marked with RFP (red) and ISCs / EBs are marked with GFP (green). **P)** Quantification of EB sphericity from the posterior midguts of fed, age-matched, mated esgReDDM > attP40 control, UAS-5HT7 or UAS-5HT7; Pbl RNAi females. **Q)** Quantification of intestinal epithelial renewal (new ECs/all ECs) of the posterior midguts from fed, age-matched, mated esgReDDM > attP40 control, UAS-5HT7 or UAS-5HT7; Pbl RNAi females. **R)** Model showing how erebotic ECs secrete serotonin to ISCs and EBs, which sense it through 5HT7, leading to EBs locating and replacing the dying ECs. P-values in (**C, E, G, M, N**) were calculated with unpaired t test with Welch correction. Adjusted *P* values in (**H-K**) were obtained in differential expression analysis of RNA-Seq with Benjamini–Hochberg correction. Normalised counts were obtained from the DESeq2 object of the respective comparison; small differences in control values between panels reflect comparison-specific size factor estimation and not different samples. P-values in (**P, Q**) were calculated with a one-way ANOVA followed by Tukey’s HSD post-hoc test for pairwise comparisons. N = biological replicates. In box plots, the center line indicates the median, the box boundaries represent the 25th and 75th percentiles, and the whiskers extend to 1.5 X IQR from the 25th and 75th percentiles. Individual dots represent separate biological replicates, except in (E, G, P), where they represent separate technical replicates (cells).

To gain better insight into the underlying mechanisms of 5HT7-mediated regulation of EB morphology, we explored RNAseq data for changes in the expression of genes involved in cytoskeletal regulation. Indeed, the expression of *pebble*, a Rho guanine exchange factor that contributes to actin cytoskeleton reorganization (Schumacher et al., 2004), and *rhophilin*, a regulator of actin stress fiber assembly (Peck et al., 2002), were deregulated upon 5HT7 misexpression (**Fig 6H-K**). To functionally test the role of actin regulation, we knocked down Pebble, which inhibited epithelial turnover (**Fig 6L, M**), but did not influence the EB numbers (**Fig 6L, N**). Moreover, the extended EB morphology as well as the elevated epithelial turnover upon 5HT7 overexpression were blunted by simultaneous Pebble knockdown (**Fig 6O-Q**). Collectively, our data are consistent with a model in which 5HT7-mediated signaling regulates EB morphology and migration, at least in part, through genes involved in cytoskeletal remodeling, and that this is necessary for epithelial turnover and integrity (**Fig 6R**).

## DISCUSSION

Our study reveals a regulatory mechanism that contributes to regional and nutrient-responsive homeostatic turnover of the *Drosophila* midgut. Firstly, we observed that turnover rates across midgut regions are much more variable than previously anticipated; in addition to areas with low or high constitutive turnover, specific areas display highly nutrient-responsive turnover. Secondly, our data show that serotonin signaling is a key regulator of turnover, thereby contributing to the regional turnover rate and allowing dying enterocytes to signal locally to promote progenitor maturation and migration. In order to do so, serotonin accumulates in ECs undergoing programmed cell death via erebosis, and it signals through the conserved 5HT7 receptor to maturing progenitor cells. Mechanistically, 5HT7 signaling controls metabolic gene expression in progenitors, promoting their maturation, and regulates progenitor migration through actin cytoskeletal regulators. Consistent with our model, defects in 5HT7 signaling impair the replacement of old ECs with newly differentiated cells, leading to slow turnover and epithelial multilayering.

Our study brings new insight into the spatial heterogeneity of intestinal turnover and its relationship with nutrient intake. In mammals, the intestinal epithelium is renewed every 4-5 days (van der Flier and Clevers, 2009). The small intestine exhibits significantly faster protein turnover than the colon (Arike et al., 2020), and while the intestinal microbiota increases turnover compared to germ-free conditions (Lesher et al., 1964), direct evidence for nutrient regulation on global intestinal turnover rate is scarce. Prior work has addressed *Drosophila* midgut turnover, reporting homeostatic turnover estimates of one week (Ohlstein and Spradling, 2006), 12 days (Jiang et al., 2009), and 21 days (Antonello et al., 2015) under normal feeding conditions. Moreover, the local turnover varies greatly between midguts of age□synchronized and co□cultured animals (Antonello et al., 2015), which may, in addition to regional differences, explain the high variation in published turnover estimates. To achieve a global view on midgut turnover regulation, we utilized quantitative organ-wide analysis with LAM (Viitanen et al., 2021). Our data revealed that the average turnover rates in fed animals display dramatic regional differences. For example, the enterocyte turnover of the R4b region had peaked already after four days of feeding, exceeding the rate of previously published observations, while R1 and R3 border region enterocytes remain very long-lived, displaying low turnover during the course of 28 days. This suggests that some enterocytes may not even be replaced throughout the animal’s lifetime, challenging the view of a universally rapid turnover of the intestinal epithelium. Previous work has shown that ISC-derived cells do not pass regional boundaries during differentiation (Marianes and Spradling, 2013), raising the possibility that the low turnover in border regions may be necessary to protect regional identities. Our results also provide evidence for a molecular determinant of regional turnover rate. Firstly, the observed intracellular serotonin accumulation is more pronounced in posterior areas, consistent with the observed high turnover rates. Secondly, inhibition of 5HT7 signaling in the progenitors strongly impaired the high turnover of the R4b region. The observation that erebotic cells accumulate high levels of serotonin, which they use to signal to progenitors, is consistent with a model that the EC state, rather than stem cell properties, serves as a key determinant of local turnover rate. Future studies are necessary to better understand the molecular mechanisms that control the rate of erebosis in the midgut epithelium. Collectively, our data show that midgut turnover is significantly more spatially heterogeneous than previously anticipated, and serotonin signaling contributes to these global differences.

In addition to the global differences in midgut turnover, our study provides new insight into the local interaction between dying ECs and their neighboring progenitors. Previous work has shown that under homeostatic conditions, healthy ECs suppress proliferation of nearby stem cells via contact inhibition. EC apoptosis initiates E-cadherin-mediated signaling in adjacent stem cells, inducing ISC division (Liang et al., 2017). However, not all ECs directly contact a stem cell, and ISC divisions generate an excess of committed progenitors even in the absence of local cell turnover (Antonello et al., 2015), indicating that soon-to-die ECs require additional mechanisms to control the maturation of local progenitors. Moreover, lineage-tracing analysis has shown that some enteroblasts migrate farther from their site of origin, a process facilitated by EMT-like formation of actin-rich protrusions (Antonello et al., 2015). How the regulation of EB maturation and migration is coordinated with the local needs of EC turnover has thus far remained unclear. Our study advances the state of the art by providing evidence that old ECs undergoing programmed cell death (erebosis), while still maintaining their position in the epithelium, accumulate high levels of serotonin to signal the need for their replacement. The identification of 5HT7 as a receiver of this signal allowed us to observe that serotonin signaling is necessary and sufficient to promote the maturation and migration-associated morphological changes of progenitors. Future efforts will be necessary to further address how replacement of erebotic epithelial cell is coordinated to avoid transient loss of barrier integrity. Our findings provide some clues about the underlying mechanisms, as we observed that serotonin-positive ECs were typically surrounded by one or more flattened EBs that formed extensive contacts with the EC. Moreover, the findings with epithelial multilayering after prolonged inhibition of enteroblast serotonin signaling suggest that EB migration to form direct contacts with the erebotic ECs may be necessary to drive the replacement of the EC from the epithelium. Such a two-step control mechanism, where 1) the dying EC activates progenitor maturation and migration and 2) the contact-forming progenitor drives the elimination of the old EC, would be a way to ensure maintenance of cellular homeostasis and epithelial integrity during turnover. Concomitantly with this study, Zipper and coworkers (Zipper et al., 2026) reported that EB guidance to the site of “worn-out” ECs is mediated through Netrin-B ligands that attract Frazzled/DCC-receptor dependent EB protrusions and subsequent EB migration in the R5 midgut region. Future studies should examine whether Serotonin and Netrin signaling cooperate to control EB migration, or whether these two signaling systems operate in parallel in distinct midgut regions.

Our study addressed the role of nutrient intake in the midgut turnover. Previous work has shown that midgut cells are highly nutrient responsive, undergoing adaptive growth in response to feeding. Nutrition increases ISC proliferation through insulin-like signaling (Choi et al., 2011; O’Brien et al., 2011). In addition, region-specific mTOR complex 1 activity increases enterocyte size and guides the fate determination of ISCs towards the absorptive lineage (Mattila et al., 2024). The highly nutrient-responsive turnover of specific midgut regions occurs in parallel with the adaptive growth control. While Insulin/mTORC1-responsive ISC proliferation and differentiation likely facilitate turnover, our findings also show that EC erebosis and serotonin accumulation depends on feeding state and correlates with turnover rate. Moreover, we observed that EC serotonin accumulation depends on the expression of Serotonin Transporter (SerT). It will be interesting to learn how enterocyte metabolism and nutrient sensing are coupled to serotonin uptake and the downstream signaling that follows. Serotonin signaling is also interconnected with EB metabolism, as our findings showed that nutrient availability is necessary for terminal differentiation even under conditions of overactivated serotonin signaling. 5HT7 activity was also shown to influence the expression of metabolic genes in EBs. Interestingly, our findings implicate the serine biosynthesis pathway as a novel metabolic driver of EB maturation. Genetic impairment of the serine biosynthesis pathway prominently suppressed EB maturation, while feeding of ectopic serine partially rescued the impaired differentiation upon inhibition of serotonin signaling. The observed role of serine biosynthesis aligns with previous observations in differentiating epidermal stem cells, whose fate determination is controlled through the serine biosynthesis pathway and its effects on alpha-ketoglutarate-mediated epigenetic regulation (Baksh et al., 2020). Future studies should address the downstream consequences of serine metabolism in EB maturation. Collectively, our findings highlight the close functional coupling between the metabolic state of both ECs and the differentiating EBs and the nutrient-responsive control of turnover by serotonin signaling.

Taken together, our findings identify serotonin as a key mediator linking nutrient availability, enterocyte state, and progenitor cell behavior to coordinate midgut epithelial renewal. By coupling the accumulation of serotonin in erebotic enterocytes to 5HT7-dependent regulation of progenitor maturation and migration, this mechanism enables local turnover demands to be translated into precise cellular replacement responses. The insights obtained from the organ-wide analyses advance the current view of midgut homeostasis by demonstrating that epithelial turnover is highly regionalized and dynamically responsive to nutritional status. By identifying the nutrient dependence of EC serotonin accumulation and the metabolic control of EB maturation downstream of 5HT7 signaling, our study highlights the key role of metabolic state as a determinant of cell fate and tissue renewal.

## MATERIALS AND METHODS

### Drosophila stocks and husbandry

Fly stocks used in this study are as follows: w; esg-Gal4, Tub-Gal80^ts^, UAS-GFP; UAS-Flp, Act>CD2>Gal4 (esgF/O) (Jiang and Edgar, 2009b), w; esg-Gal4, Tub-Gal80^ts^, UAS-GFP (esg-Gal4^ts^) (Jiang and Edgar, 2009a), w; MyO1A-Gal4, Tub-Gal80^ts^, UAS-GFP (MyO1A-Gal4^ts^) (Jiang et al., 2009), w; esg-GAL4, UAS-mCD8-GFP; tub-G80^ts^, UAS-H2B-RFP (esgReDDM) (Antonello et al., 2015), w; esg-GAL4, UAS-CFP, Su(H)-GFP, His-RFP (Cell fate sensor) (Martin et al., 2018), 5HT7-RNAi (Bloomington 27273), Vmat-RNAi (Bloomington 44471), SerT-RNAi (VDRC v100584), Aay RNAi (Bloomington 28577), Pbl-RNAi (Bloomington 28343), Luciferase-RNAi (Trip control Bloomington 31603), UAS-5HT7 (attP40 site, this study), attP40 control line (Bloomington 25709). Flies were maintained at 25°C on a standard laboratory diet containing agar 0.6% (w/v), malt 6.5% (w/v), semolina 3.2% (w/v), baker’s yeast 1.8% (w/v), nipagin 2.4%, and propionic acid 0.7%. In experiments using the temperature-sensitive Gal80^ts^, flies were reared at +18°C and then transferred to +29°C to inactivate the Gal80^ts^ protein, allowing UAS-Gal4-driven transgene expression. The exact timings of the dietary treatments and temperature switch in each experiment are provided in the figure legends.

### Dietary treatments

The fasting diet consisted of 2% sucrose (w/v) in a medium containing 0.5% agar (w/v), nipagin 2.4%, propionic acid 0.7%, and NaOH to adjust the pH to 6.8. For L-Tryptophan (Sigma 51145) and 5HTP (Sigma H9772) feeding experiments, L-Tryptophan and 5HTP were added to the fasting medium at the concentrations of 25 mM and 7.5 mM, respectively, unless otherwise mentioned. Flies were kept on a chemically defined (holidic) diet (Piper et al., 2014), referred to as the fed condition.

### Immunohistochemistry

For immunofluorescence staining, midguts were dissected in phosphate-buffered saline (PBS) and fixed in 4-8% paraformaldehyde for at least 2 hours. Samples were washed with 0.1% Triton X-100 in PBS (PBT) and blocked in 1% bovine serum albumin for at least 30 minutes. Subsequently, intestines were stained with anti-5HT (1:1000) (Sigma S5545), anti-ANCE (1:1000) (Kerafast ELD038), anti-Horseradish Peroxidase HRP 1:200 (Jackson ImmunoResearch Laboratories 2338967). The samples were then washed again with PBT and stained with secondary antibodies for at least an hour. Tissues were then washed and mounted in Vectashield mounting medium with DAPI (Vector Laboratories).

### Ex vivo imaging of midguts

Ex vivo imaging was performed as described previously (Marchetti et al., 2022) with a few modifications. M3 media was used instead of Schneider’s media for the dissection of midguts, fetal bovine serum was used instead of bovine serum albumin in all steps, and penicillin/streptomycin mixture was used as the antibiotic. In short, midguts were dissected in M3 media and then incubated with 10 µM Hoechst. Midguts were then embedded in a 1% low-melting agarose gel in M3 medium, and an imaging medium based on fly extract was added. Imaging began within 20 minutes of dissection using the Aurox clarity spinning-disc confocal microscope.

### Microscopy and image processing

Fixed midguts were mounted between a microscope slide and a coverslip with 0.12 μm spacers. Samples were imaged using the Aurox Clarity spinning disc confocal microscopes, the SP8 confocal microscope, the Andor Dragonfly 505 high-speed spinning disc confocal or the Stellaris 8 confocal microscope. For regionally defined cellular phenotyping (LAM), images were processed with ImageJ and segmented with StarDist as previously described (Viitanen et al., 2021). Intensities were measured as the average voxel or pixel intensity within the segmented regions of interest. Cell sphericity was calculated using Imaris and manual segmentation of cell borders.

### FACS and RNA sequencing

ISCs and EBs expressing GFP (esgGAL4^ts^ > UAS-GFP) were dissociated for FACS-sorting as described previously (Dutta et al., 2015). Approximately 80 posterior midguts per sample were dissected in DPBS, then transferred to a DNAlow-bind (Eppendorf) pre-coated tube containing filtered 1% BSA in PBS. Midguts were dissociated in 1 mL of 10x TrypLE (Thermo Fisher Scientific) at 25°C at 800rpm for 30 minutes. After complete dissociation, cells were centrifuged at 600 × g for 10 minutes at 4 °C. The pellet was resuspended in 500 µl of filtered 1% BSA in PBS and then filtered through an equilibrated 40 µm cell strainer. Propidium iodide was added to the samples at a final concentration of 2 µg/ml to distinguish nonviable cells. Cells were sorted using Aria Fusion flow cytometer (BD Biosciences) based on GFP. RNA was isolated using the Arcturus PicoPure RNA isolation kit (Thermo Fisher Scientific) and subsequently amplified using the RiboAmp HS-Plus kit (Thermo Fisher Scientific). Libraries were generated using the TruSeq Stranded mRNA Library Prep Kit (Illumina) and subsequently sequenced as 50 base-pair single-end reads on Illumina HiSeq2500.

RNA-seq reads were processed with the nf-core/rnaseq pipeline (version 3.18.0) (Ewels et al., 2020) using Nextflow (version 24.10.3) and Singularity containers. Ribosomal RNA reads were removed, and adapters and low-quality bases were trimmed with Trim Galore (version 0.6.10). Reads were aligned to the *Drosophila melanogaster* reference genome (FlyBase release r6.54) using STAR (version 2.7.11b), and transcript abundance was quantified with Salmon (version 1.10.3). Salmon-derived transcript counts were summarized to gene-level counts using tximport (version 1.32.0). The raw and processed sequencing data is deposited into the GEO repository (GSE335701).

Differential expression analysis was performed in R/Bioconductor using DESeq2 (version 1.44.0). Because the experiment comprised two comparisons sharing a common control group, differential expression was analysed as two separate DESeq2 datasets, control versus overexpression and control versus knockdown, each containing only the libraries relevant to that comparison. The identical control libraries were used in both analyses. As size factors, dispersion estimates and independent filtering are computed from the set of samples present in a given DESeq2 object, the normalised counts obtained for the shared control libraries differ slightly between the two analyses. Normalised counts shown in figures therefore correspond to the analysis in which that comparison was performed and are not directly comparable across the two comparisons. Genes with fewer than 10 estimated counts in at least three of four biological replicates in at least one experimental group were excluded before analysis. P-values were adjusted for multiple testing using the Benjamini–Hochberg method. Differentially expressed genes were subjected to over-representation analysis using clusterProfiler (version 4.12.6) with Gene Ontology, KEGG, Reactome, and WikiPathways annotations. Enrichment p-values were corrected using the Benjamini–Hochberg method.

### Isolation of mouse small intestinal crypts and organoid culture

Small intestinal crypts were isolated from female C57BL/6JRccHsd mice (wild-type or Lgr5-EGFP-IRES-creERT2) as previously described (Andersson et al., 2025). To isolate the crypts, the small intestine was first flushed with ice-cold PBS and opened longitudinally to remove the mucus. After being cut into small fragments, the tissue was incubated on ice for 1 hour and 45 minutes with four changes of 10 mM EDTA in PBS. During the first three 15-minute incubations, the villus fractions were discarded after each manual shaking. After a final 60-minute incubation, the crypt fraction was collected by vigorous manual shaking. The collected crypts were then filtered through a 70 µm nylon mesh, centrifuged, and washed with cold PBS to remove any residual EDTA. Isolated crypts were kept on ice and resuspended in Advanced DMEM/F12 (Gibco). For culture, crypts were plated in 20 µL plugs of 60% Matrigel (Corning) at the center of 48-well plates. The culture medium, referred to as ENR medium, consisted of Advanced DMEM/F12 (Gibco) supplemented with 1× GlutaMAX (Gibco), 100 U/mL penicillin and streptomycin, 10 mM HEPES, 1× B-27 supplement (Gibco), 1× N-2 supplement (Gibco), 50 ng/mL mouse EGF (R&D Systems), 100 ng/mL mouse noggin (PeproTech), and 500 ng/mL mouse R-spondin1 (R&D Systems). For the first two days of culture, 10 µM of Y-27632 was added to the medium. The culture medium was replaced every 2-3 days. The Htr7 agonist LP-211 (Sigma-Aldrich) was dissolved in DMSO to 100 µM and added to the organoid culture immediately after crypt isolation. Organoid budding was counted from brightfield images taken on day 6 or 7. Four to seven images were captured per biological replicate, with each image containing 40 to 110 organoids. The data presented are from four independent experiments using crypts isolated from four different mice.

### Statistical analysis

Statistical analyses were performed in R/Bioconductor. A two-tailed or one-tailed t test, a paired sample t test, or a two-way analysis of variance (ANOVA) in conjunction with Tukey’s post hoc test was used. P values for the RNA sequencing experiment were calculated using the Wald significance test and adjusted with Benjamini and Hochberg correction for multiple testing. The exact test used in each experiment is indicated in the figure legend.

## Supporting information

Supplementary figures_Lamichane

## RESOURCE AVAILABILITY

### Lead contact

Further information and requests for resources and reagents should be directed to the lead contact, Ville Hietakangas.

## AUTHOR CONTRIBUTIONS

Conceptualization, N.L., J.M. and V.H.; investigation, N.L., P.B., T.R., K.G., M.R., C.B., S.A., J.B. and O.K.; formal analysis, N.L., P.B., T.R., K.G., M.R., O.D., T.K., A.V. and C.B.; writing – Original draft, N.L., J.M. and V.H.; funding acquisition, V.H., J.M. and N.L.; resources, P.K., V.H.; supervision, J.M. and V.H.

## ACKNOWLEDGEMENTS

We thank Bloomington *Drosophila* Stock Center, Lucy O’Brien and Maria Dominquez for providing fly stocks. We thank Cise Kizilirmak and Ella Salminen from Pekka Katajisto’s lab for their kind help in setting up the methods for the small intestinal organoid experiment. This work was supported by the Research Council of Finland (# 336197, 372024 and 374086 to V.H.), Sigrid Jusélius Foundation (to V.H. and J.M.), Novo Nordisk Foundation: NNF19OC0057478, NNF22OC0078419 and NNF25OC0105265 (to V.H.), Jane & Aatos Erkko Foundation (to J.M.), and the Integrative Life Science Doctoral Program (to N.L.). This study was facilitated by the University of Helsinki *Drosophila* core facility (Hi-Fly), DNA Sequencing and Genomics Laboratory (supported by HiLIFE and Biocenter Finland funding), the Light Microscopy Unit (LMU), supported by Biocenter Finland, the Biomedicum Imaging Unit (BIU), and the Helsinki Institute of Life Science.

