## Supplementary figures_Lamichane for "Nutrient-responsive midgut turnover is spatially controlled by serotonin signaling"

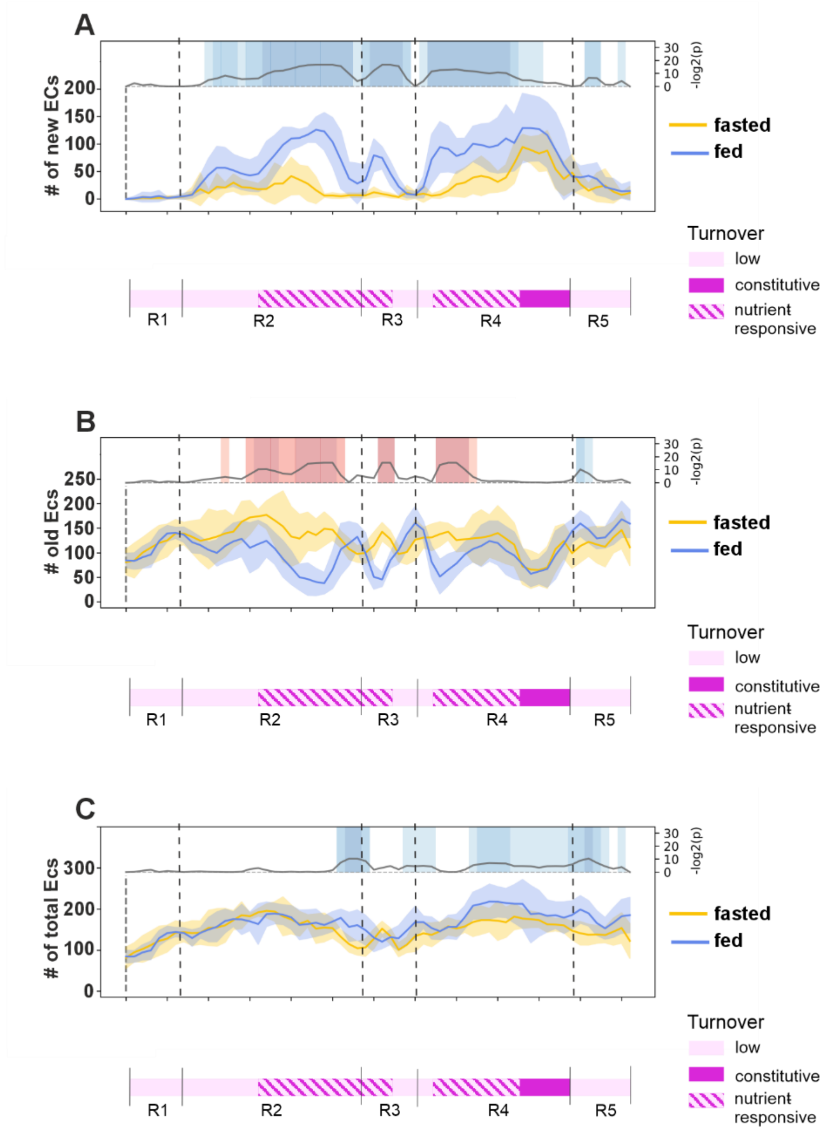


**Figure S1.**

**A)** New ECs are formed in the nutrient-responsive regions. LAM plot showing the amount of newly formed ECs (polyploid RFP+ GFP- cells) along the midgut of age-matched mated esgReDDM > luc RNAi fed or fasted females. Standard deviation is marked by line shading**. B)** Old ECs are replaced in the nutrient-responsive regions. LAM plot showing the number of old ECs (polyploid RFP- GFP- cells) along the midgut of age-matched mated esgReDDM > luc RNAi fed or fasted females. Standard deviation is marked by line shading. **C)** Overall EC numbers only slightly increased in the posterior midgut. LAM plot showing the number of ECs (polyploid GFP- cells) along the midgut of age-matched mated esgReDDM > luc RNAi fed or fasted females. Standard deviation is marked by line shading. The schemes below each graph display the turnover patterns of the midgut regions. p values are calculated by Mann-Whitney-Wilcoxon U test using a 3-bin window.

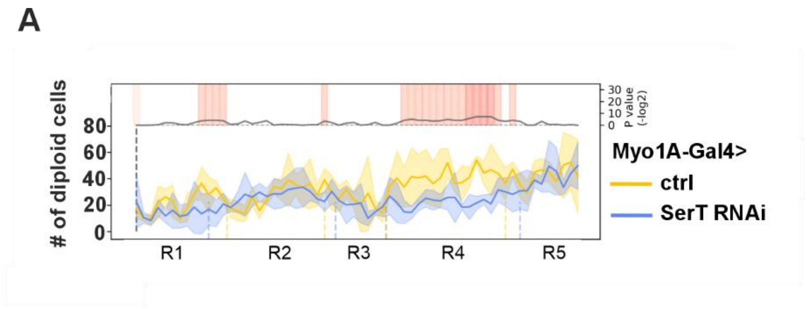


**Figure S2.**

**A)** SerT in ECs regulates stem cell activity. Quantification of small cells (mostly ISCs and EBs) along the midgut of fed, age-matched, mated Myo-GAL4 > luc RNAi or SerT RNAi females. Standard deviation is marked by line shading. p values are calculated by Mann-Whitney-Wilcoxon U test using a 3-bin window.

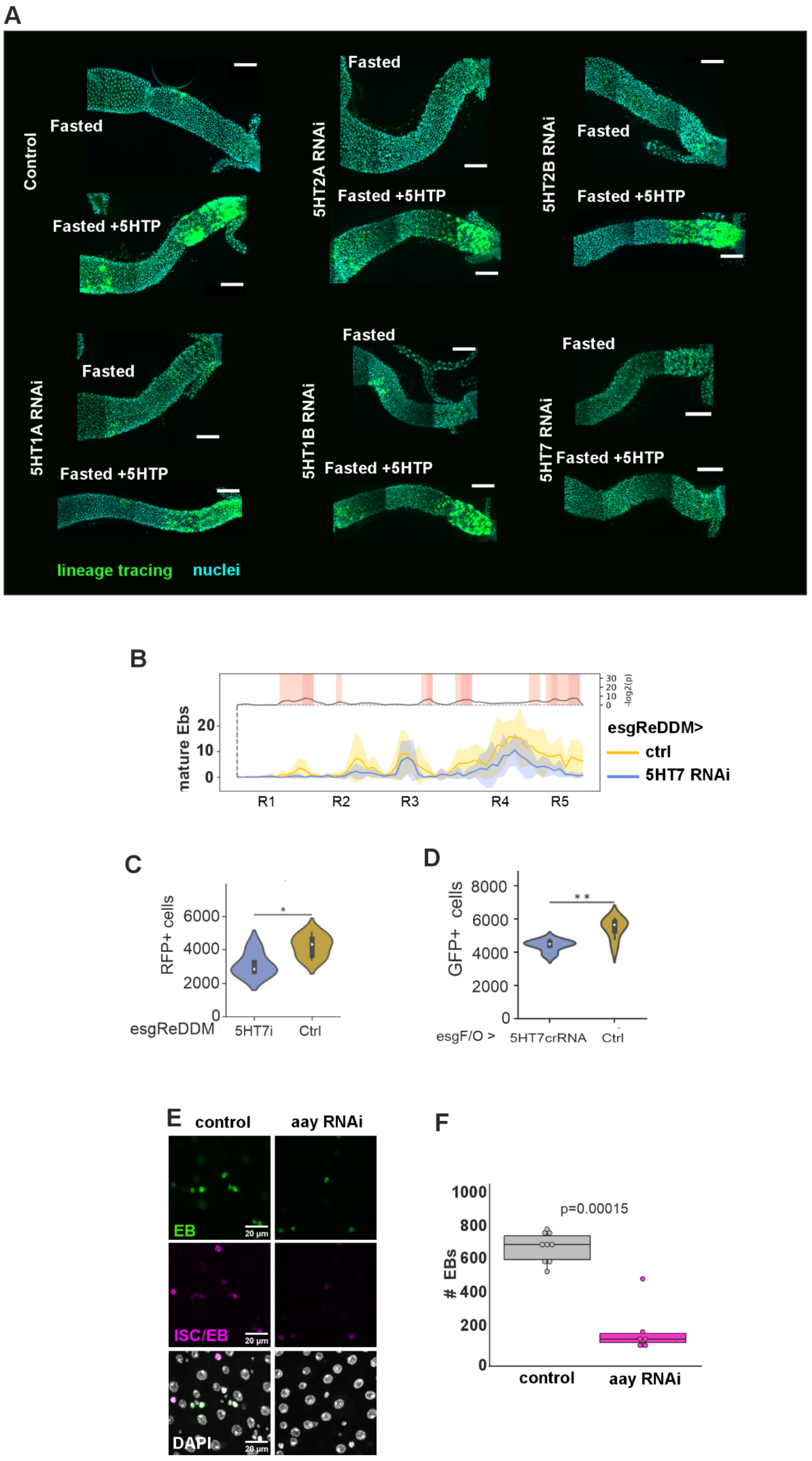
A
Control
Fasted
Fasted +5HTP
5HT2A RNAi
Fasted
Fasted +5HTP
5HT2B RNAi
Fasted
Fasted +5HTP
5HT1A RNAi
Fasted
Fasted +5HTP
5HT1B RNAi
Fasted
Fasted +5HTP
5HT7 RNAi
Fasted
Fasted +5HTP
lineage tracing
nuclei
B
mature Ebs
20
10
0
30
20
10
0
-log2(p)
R1
R2
R3
R4
R5
esgReDDM>
ctrl
5HT7 RNAi
C
RFP+ cells
6000
4000
2000
0
*
esgReDDM
5HT7i
Ctrl
D
GFP+ cells
8000
6000
4000
2000
0
* *
esgF/O >
5HT7crRNA
Ctrl
E
control
aay RNAi
EB
ISC/EB
DAPI
20 µm
F
### EBs
1000
800
600
400
200
0
p=0.00015
control
aay RNAi

**Figure S3.**

**A)** 5HT7 expression in ISCs and EBs is necessary for the midgut response to 5HTP feeding. Representative image of age-matched, mated esgF/O > luc ctrl, 5HT1A, 5HT1B, 5HT2A, 5HT2B or 5HT7 RNAi posterior midgut fed with 7.5 mM 5HTP. **B)** 5HT7 regulates maturation of EBs. Full midgut regional quantification of mature EBs of fed, age-matched, mated esgReDDM > luc RNAi or 5HT7 RNAi females. Standard deviation is marked by line shading. **C)** Quantification of RFP-positive cells (stem cell clones) in full midguts of age-matched, mated esgReDDM > luc RNAi (control) or 5HT7 RNAi females fed with 7.5 mM 5HTP. **D)** Quantification of GFP-positive cells (stem cell clones) in full midguts of fed, age-matched, mated esgF/O > pCFD8, UAS-Cas12 (control) or 5HT7 crRNA, UAS-Cas12 females. **E)** Representative images of ISCs (CFP+ and GFP-) and EBs (CFP+ and GFP+) from posterior midguts of fed, age-matched, mated Cfs-GAL4 > luc RNAi or Aay RNAi females. **F)** Quantification of EB numbers from the experiment depicted in E. P-values in (**B**) are calculated by Mann-Whitney-Wilcoxon U test using a 3-bin window. P-values in (**C-F**) were calculated with unpaired t test with Welch correction. In box plots, the center line represents the median, the box boundaries represent the 25th and 75th percentiles, and the whiskers extend to 1.5 X IQR from the 25th and 75th percentiles. Individual dots represent separate biological replicates.

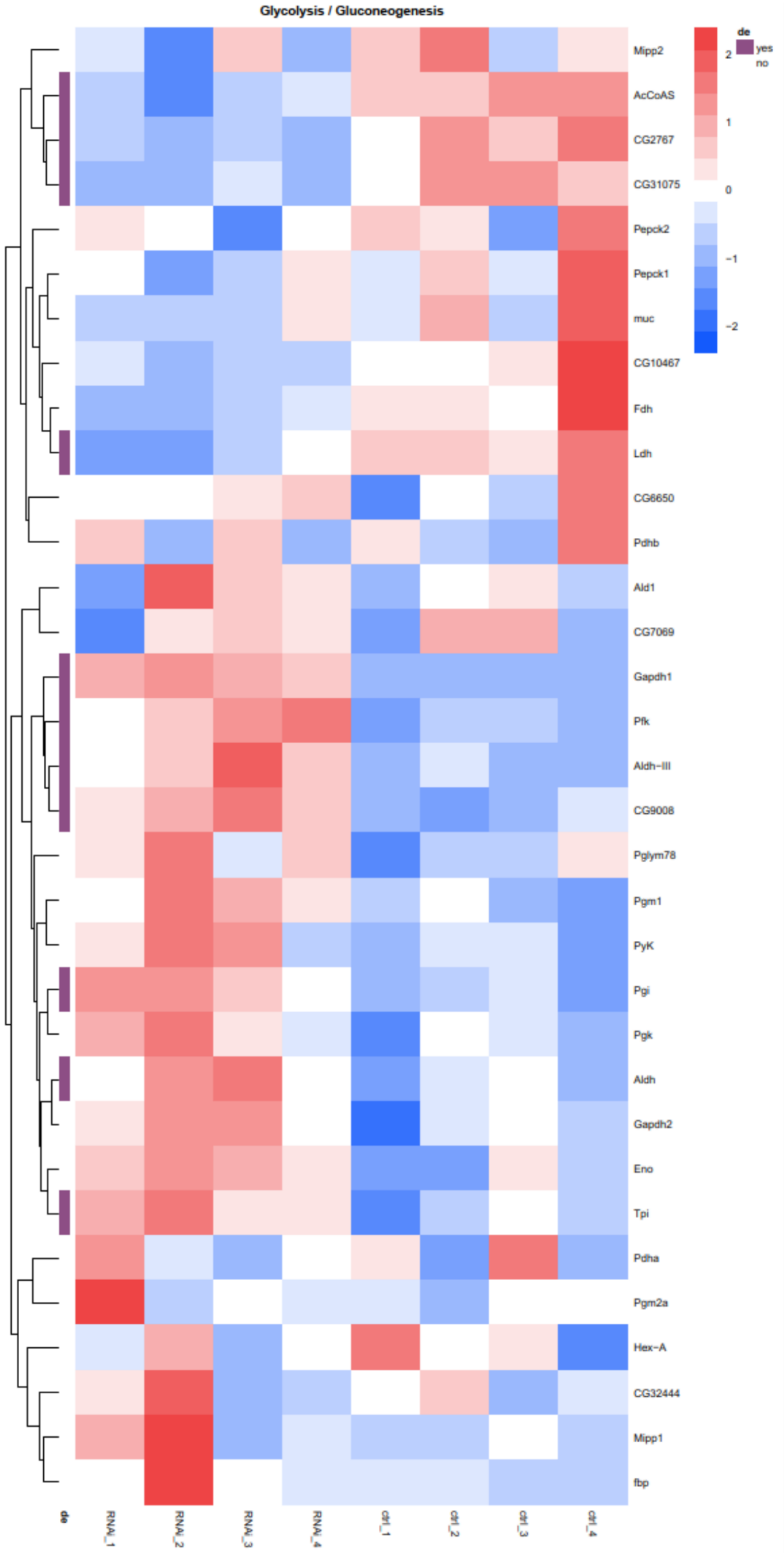
Glycolysis / Gluconeogenesis
Mipp2
AcCoAS
CG2767
CG31075
Pepck2
Pepck1
muc
CG10467
Fdh
Ldh
CG6650
Pdhb
Ald1
CG7069
Gapdh1
Pfk
Aldh−III
CG9008
Pglym78
Pgm1
PyK
Pgi
Pgk
Aldh
Gapdh2
Eno
Tpi
Pdha
Pgm2a
Hex−A
CG32444
Mipp1
fbp
de
RNAi_1
RNAi_2
RNAi_3
RNAi_4
ctrl_1
ctrl_2
ctrl_3
ctrl_4
2
1
0
−1
−2
de
yes
no

**Figure S4.**

Heatmap of enriched metabolic pathway Glycolysis/Gluconeogenesis showing differentially expressed genes between control and 5HT7 knockdown samples.
